# Changing color and intensity of LED lighting across the day impacts on human circadian physiology, sleep, visual comfort and cognitive performance

**DOI:** 10.1101/2020.04.21.771832

**Authors:** O. Stefani, M. Freyburger, S. Veitz, T. Basishvili, M. Meyer, J. Weibel, K. Kobayashi, Y. Shirakawa, C. Cajochen

## Abstract

We examined whether dynamic light across a scheduled 16-h waking day influences cognitive performance, visual comfort, melatonin secretion, sleepiness and sleep under strictly controlled laboratory conditions of 49-h duration.

Participants spent the first 5-h in the evening under standard lighting, followed by an 8-h nocturnal sleep episode at habitual bedtimes. Thereafter volunteers either woke up with static daylight LED (100 lux and 4000 Kelvin) or with a dynamic daylight LED that changed color (2700 – 5000 Kelvin) and intensity (0 - 100 lux) across the scheduled 16-h waking day. This was followed by an 8-h nocturnal treatment sleep episode at habitual bedtimes. Thereafter, volunteers spent another 12-h either under static or dynamic light during scheduled wakefulness.

Under dynamic light, evening melatonin levels were less suppressed 1.5hours prior to usual bedtime, and participants felt less vigilant in the evening compared to static light. Sleep latency was significantly shorter in both the baseline and treatment night compared to the static light condition while sleep structure, sleep quality, cognitive performance and visual comfort did not significantly change. Our results support the recommendation of using blue-depleted light and low illuminances in the late evening, which can be achieved by a dynamically changing daylight LED solution.

## 1. Introduction

In our modern societies we spend more and more time indoors under artificial light ^1^. As a consequence, we expose ourselves to less sunlight during the day and more artificial (electric) light, particularly in the evening after sunset potentially delaying our internal clock with the risk of desynchronizing endogenous rhythmicity and external time demands (i.e. social/working schedule). This is typically associated with difficulties falling asleep at night and getting up in the morning without an alarm clock. In order to avoid such circadian misalignments, it is recommended to increase Zeitgeber (i.e. “time giver”) strength by increasing light exposure during the day and avoid light at night. The wrong light at the wrong time of day is also associated with negative effects on well-being and sleep ^2–5^. Thus, a prerequisite for good sleep and health are internal biological clocks that are well synchronized with the 24-hour day on earth with its natural light-dark changes. Internal clocks are kept in sync by signals from a part of the brain that responds to changes in light and darkness. Intrinsically photosensitive retinal ganglion cells (ipRGC) use the photopigment melanopsin to transmit light stimuli via the retinohypothalamic tract into different brain areas, primarily to our internal “master clock”, the nucleus suprachiasmaticus (SCN) ^6–9^. Melanopsin is very sensitive to short-wave radiation, i.e. blue light, making light of these wavelengths highly effective in their function as a Zeitgeber ^10,11^. Especially blue light and white light with a high proportion of short-wave radiation during the evening and at night counteracts the natural increase in sleepiness and suppresses endogenous melatonin secretion ^12,13^. Melatonin, sometimes referred to as the “dark hormone” is secreted periodically and is used as a robust marker for the state of the circadian system ^14^. Light has been shown to have acute and delayed effects on this rhythm. The production of melatonin is acutely inhibited by light ^14,15^. Brainard et al. ^13^ and Thapan et al. ^12^ measured the strongest melatonin suppression with monochromatic light at a wavelength of 464 nm. Delayed effects of light are typically measured in so-called phase response protocols, where lights’ Zeitgeber strength on circadian melatonin or core body temperature phase were quantified i.e. non-parametric effect of light ^16,17^. Also for light’s phase shifting capacity a short-wavelength dominance has been reported ^18^. These effects of light exposure are not limited to strong illumination ^19^ and have also been observed when comparing different usage of home lighting in a field study ^20^.

So-called Human Centric Lighting (HCL) aims at incorporating the above mentioned non-visual effects of light by dynamically changing correlated color temperature (CCT) and illuminance across the day, thereby altering short-wave radiation. Thus, HCL aims at influencing non-visual forming effects to support circadian physiology and good sleep in humans. In animal studies, simulated dawns enhanced circadian entrainment (the match of rhythmic physiological events to environmental oscillations, such as the natural light-dark cycle, with the result that both oscillations have the same frequency). in comparison to abrupt light transitions ^21–24^. These results suggest that natural twilight simulation increases the Zeitgeber strength of the light-dark (LD) cycles through parametric rather than non-parametric entrainment mechanisms (i.e. parametric entrainment is considered as a continuous process in which the circadian oscillator constantly accelerates and decelerates to adapt to the environment, while the circadian oscillator is advanced or delayed every day in an almost instantaneous manner in a non-parametric view of entrainment ^25^. Effects of a simulated dawn on circadian rhythms in humans was investigated by Danilenko et al. ^26^ with a halogen lamp, which delivered an intensity change from 0.001 to 1000 lux. The rate of change of illuminance represented a natural sunrise. Control participants remained under dim light conditions with an alternating light dark cycle (< 30:0 lux). Their results showed that a replacement of the last 1.5 h of darkness by a natural dawn stimulus (averaging 155 lux) was sufficient to maintain an entrained phase position in comparison to the control situation. In a study in which both the color temperature (1090 - 2750 K) and illuminance at the eye (0 – 250 lux) was changed during wake up in the morning, mood, well-being and cognitive performance increased ^27,28^, and the cardiac control during the awakening process was better ^29^ in comparison to 8 lux dim light together with a technicians voice as a wake up signal. A twilight simulation was also found to be antidepressant ^30^ and to reduce sleep disorders ^31^. In a study comparing dynamic light with static light for office workers illuminance was changed between 500 and 700 lux and the CCT between 3000 and 4700 K. No positive effects on sleep, vitality, headaches and productivity of dynamic light were found. However, the employees were subjectively more satisfied with the dynamic light than with the static light ^32^. To date, non-visual light effects were predominantly studied during the evening or at night, and to a lesser extent during the day with inconclusive findings. Twilight studies and the combination of the three time points (i.e. night, day and twilight) investigating the continuous change of light during 16 hours of wakefulness are rare, and therefore it remains unclear if these changes improve circadian physiology, visual comfort, cognitive performance, and sleep.

In the present study, we compared a static lighting condition “sLED” (4000 K) with a dynamically changing light “dynLED”, that started in the morning with 3500 K/< 1 lux incrementally increasing until reaching 5000 K/100 lux at 10 am. CCT and illuminance continuously decreased in the afternoon finally reaching 2700 K/< 1 lux at bedtime in the dynamic condition. This light profile is often named “circadian” or “human centric” lighting since it is generally claimed to have various, sometimes unspecified positive effects on humans, which have not yet been tested rigorously. We expected less melatonin suppression in the evening prior bedtime and better sleep, as indexed by more EEG delta activity in the treatment night, after the dynamic in comparison to the static light condition. Furthermore, visual comfort, alertness and cognitive performance during the 16 h of wakefulness during the dynamic light condition is improved compared to static light during times when exposed to 5000 K compared to static light at 4000 K. In the evening however, we expected better cognitive performance and higher alertness during static light condition due to both higher intensity and higher correlated color temperature.

## 2. Results

### 2.1. Cognitive Performance

#### 2.1.1. Psychomotor Vigilance Task (PVT)

The time course of PVT performance was rather stable across the entire day during both light conditions and did not reveal any significant main effect of the factor ‘light’ for the different measures [i.e. median reaction time (RT), lapses, the 10% fastest and 10% slowest RTs]. Also, the factor ‘time of day’ and the interaction ‘light’ x ‘time of day’ did not yield any statistically significant effects, neither for median RT, the 10% slowest or 10% fastest RTs nor the attentional lapses.

#### 2.1.2. Working Memory Performance (n-back)

There was no effect of the factor ‘light’. The factor ‘time of day’ yielded significance with better performance in the course of the scheduled waking day (F_6,62_ = 2.66, *p* = 0.023). The interaction ‘light’ x ‘time of day’ was not statistically significant.

### 2.2. Subjective Variables

#### 2.2.1. Visual Comfort

Participants rated visual comfort (i.e. the combined items brightness and CCT) similar for both light conditions. Also, the factor ‘time of day’ and the interaction ‘light’ x ‘time of day’ were not statistically significant.

#### 2.2.2. Brightness

There was neither an effect of the factor ‘light’ nor any significant effect for the factor ‘time of day’ but the interaction term ‘light’ x ‘time of day’ was statistically significant (F_7,92_ = 3.76, *p* = 0.001). *Post-hoc* tests revealed that participants rated brightness better at 11 pm during sLED compared to dynLED (t = 4.31, *p* = 0.004; Figure 1a).

**Figure 1:**
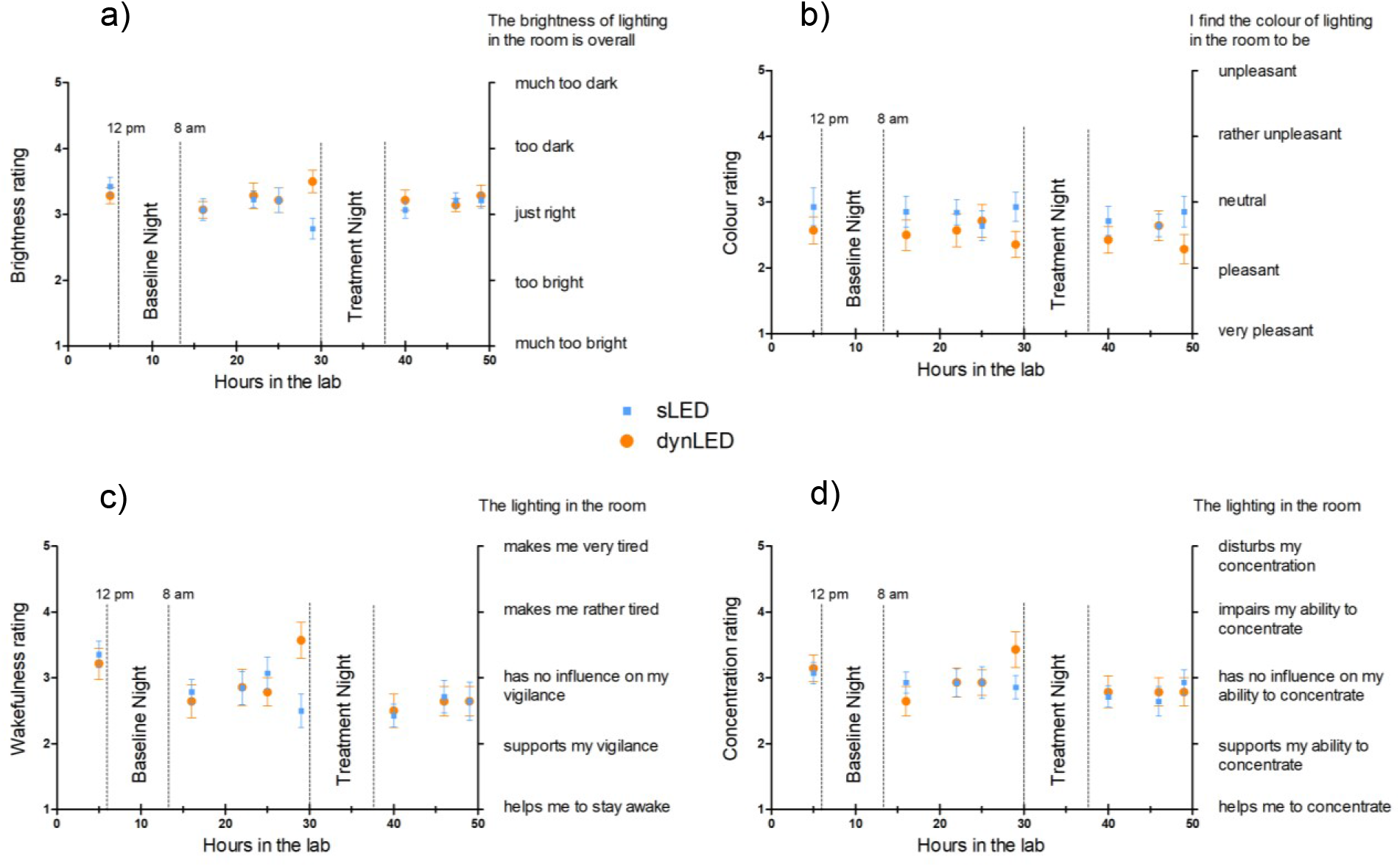
Time course of diurnal subjectively rated brightness (a), colour (b), wakefulness (c) and concentration (d) for sLED and dynLED on 5-Point Likert-Scales plotted against time in hours spent in the lab. Depicted are the means and standard errors of the mean (n=14). Brightness was rated better at 11 pm during sLED compared to dynLED (t = 4.31, *p* = 0.004) (a). Participants tended to rate colour better for dynLED (factor light: F_1,26_ = 3.57, p = 0.07) than sLED (b). Participants were feeling less vigilant during the evening (F_7,79_ = 3.81, *p* = 0.001) (c). The factor ‘time of day’ showed a significant effect with participants being less concentrated in the evening in general (F_7,81_= 2.29, *p* = 0.035) (d).

#### 2.2.3. Correlated Color Temperature (CCT)

Overall, participants tended to rate CCT better for dynLED (factor light: F_1,26_ = 3.57, *p* = 0.07) than sLED. There was neither a significant main effect of ‘time of day’ nor a significant interaction of the factors ‘light’ and ‘time of day’ (Figure 1b).

#### 2.2.4. Perception of Vigilance

There was no significant main effect for the factor ‘light’. The factor ‘time of day’ was significant (F_7,79_ = 3.81, *p* = 0.001) with participants feeling less vigilant during the evening. In addition, the interaction term ‘light’ x ‘time of day’ yielded significance (F_7,97_ = 4.76, *p* = 0.0002). *Post-hoc* comparisons indicated that the volunteers felt less vigilant under dynLED compared to sLED in the evening at 11 pm (t = 3.36, *p* = 0.035; Figure 1c).

#### 2.2.5. Perception of Concentration

The factor ‘light’ did not yield significance, while the factor ‘time of day’ showed a significant effect with participants being less concentrated in the evening in general (F_7,81_ = 2.29, *p* = 0.035; Figure 1d). The interaction term ‘light’ x ‘time of day’ yielded no significance.

#### 2.2.6. Subjective Sleepiness

Subjective sleepiness rated on the Karolinska Sleepiness Scale did not significantly differ between sLED and dynLED (data not shown). It showed, however a typical diurnal profile with lower sleepiness during the day and increased sleepiness in the evening (factor ‘time of day’: F_40,496_ = 5.69, *p* < 0.001). The interaction term ‘light’ x ‘time of day’ yielded no significance.

**Table 1:** Results of the analysis of variance for different subjective variables and cognitive performance over the time course of the study. In bold results with *p* < 0.05.

| Analysis of variance |  |  |  |
| --- | --- | --- | --- |
|  | Light | Time of day | Light*Time of day |
| *Cognitive Performance* |  |  |  |
| PVT | $F_{1,32} = 1.79, p = 0.19$ | $F_{7,72} = 0.32, p = 0.94$ | $F_{7,91} = 0.50, p = 0.83$ |
| n-back | $F_{1,25} = 0.37, p = 0.55$ | **$F_{6,62} = 2.66, p = 0.023$** | $F_{6,71} = 0.72, p = 0.64$ |
| *Subjective Variables* |  |  |  |
| Visual comfort | $F_{1,25} = 0.88, p = 0.36$ | $F_{7,82} = 0.63, p = 0.73$ | $F_{7,93} = 0.49, p = 0.84$ |
| Brightness | $F_{1,30} = 0.87, p = 0.36$ | $F_{7,83} = 1.19, p = 0.32$ | **$F_{7,92} = 3.76, p = 0.001$** |
| CCT | $F_{1,26} = 3.57, p = 0.07$ | $F_{6,71} = 0.72, p = 0.64$ | $F_{7,95} = 1.59, p = 0.15$ |
| Vigilance | $F_{1,23} = 0.08, p = 0.78$ | **$F_{7,79} = 3.81, p = 0.001$** | **$F_{7,97} = 4.76, p = 0.0002$** |
| Concentration | $F_{1,24} = 0.06, p = 0.81$ | **$F_{7,81} = 2.29, p = 0.035$** | $F_{7,97} = 1.65, p = 0.1317$ |
| Sleepiness | $F_{1,88} = 0.62, p = 0.43$ | **$F_{40,496} = 5.69, p < 0.001$** | $F_{40,504} = 1.06, p < 0.37$ |

### 2.3. Melatonin

#### 2.3.1. Diurnal Melatonin Profile

Salivary melatonin exhibited the typical diurnal profile with lower levels during daytime and increasing levels in the evening (factor ‘time of day’: F_41,441_ = 8.01, *p* < 0.0001, Figure 2). There was no significant effect of the factor ‘light’ on the diurnal melatonin profile (F_1,85_ = 0.33, *p* = 0.57), while it tended to interact with the factor “time of day” (F_41,460_ = 1.33, *p* = 0.089). *Post-hoc* comparisons revealed significantly higher melatonin levels under dynLED compared to sLED 1.5 h prior to bedtime (t = 2.11, *p* = 0.035) (Figure 2).

**Figure 2:**
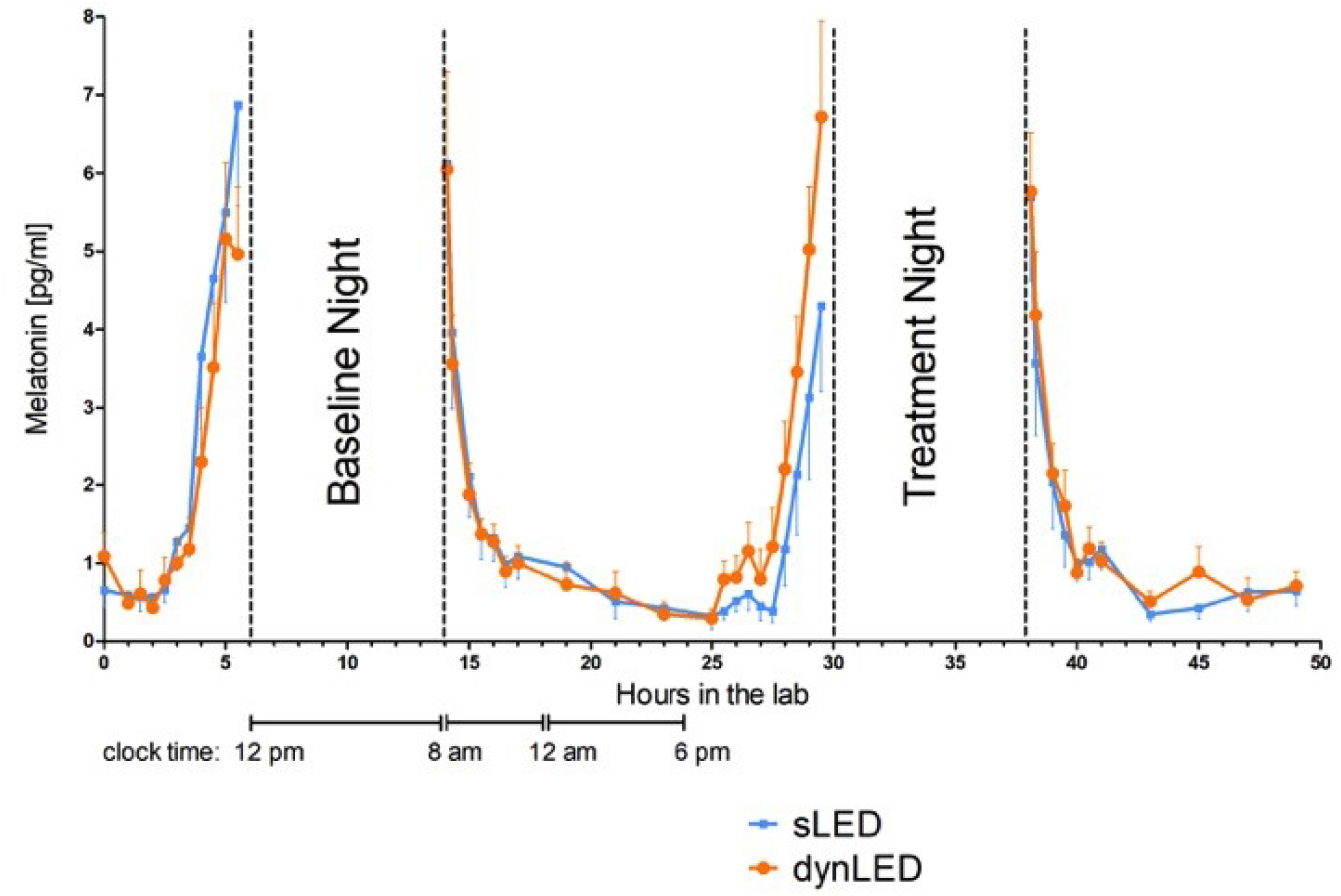
Time course of diurnal salivary melatonin profiles during the experimental conditions, sLED (blue) and dynLED (orange) in pg/ml plotted against time in hours spent in the lab (and clock time i.e. average clock times according to the participant’s habitual bedtimes). Depicted are the average melatonin levels across participants (mean values, n=14; ±SEM) and the standard errors of the mean.

#### 2.3.2. DLMO (Dim Light Melatonin Onset)

The factor ‘light’ did not show significant differences between sLED and dynLED. The factor ‘night’ and the interaction term ‘light’ x ‘night’ was statistically significant (F_1,12_ = 12.77, *p* = 0.004) and (F_1,7_ = 16.69, *p* = 0.005) respectively. *Post-hoc* tests revealed a significantly earlier (54 minutes) DLMO during the baseline evening (2.800 K) than during the treatment evening (4.000 K) in sLED condition (t = 4.85, *p* = 0.008). Such a delay in the DLMO between the baseline evening (2800 K) and the treatment evening (2700 K) was not present in the dynLED condition (9 minutes earlier DLMO during the baseline compared to treatment evening, non-significant) (Figure 3).

**Figure 3:**
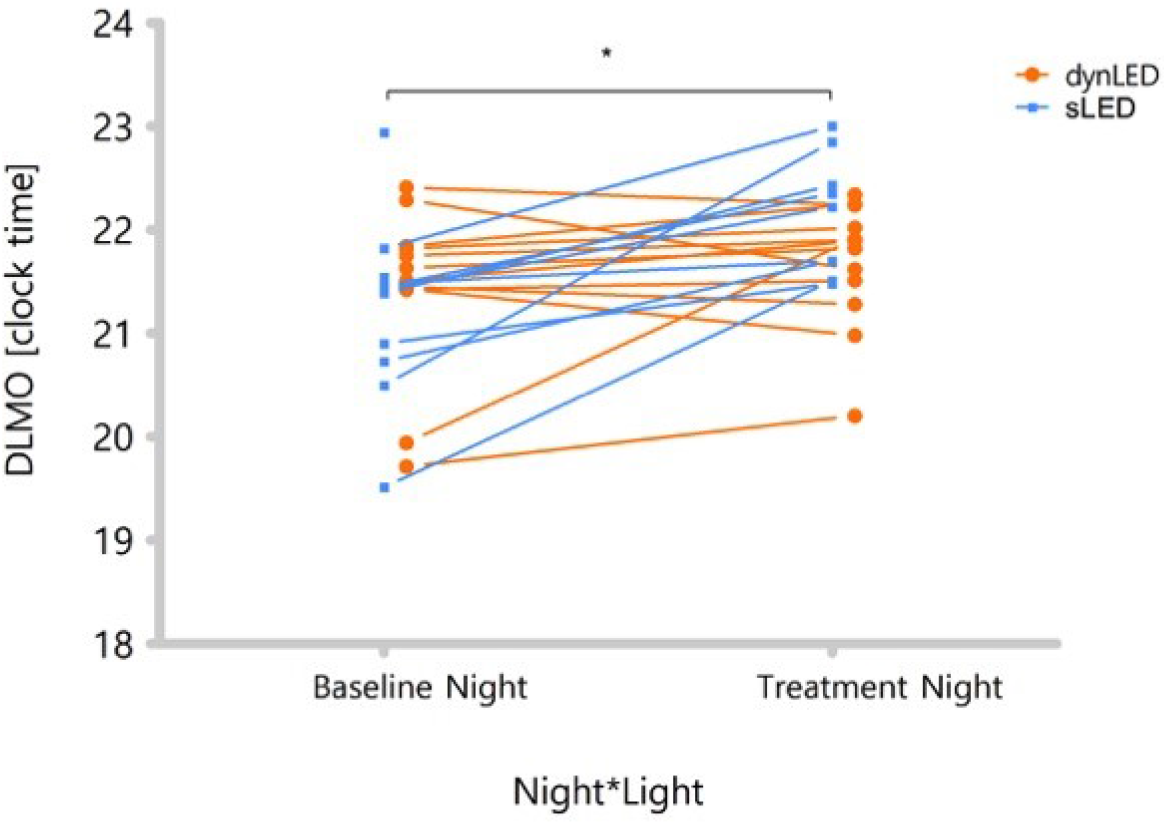
Dim light melatonin onset time prior to the baseline and treatment night under sLED (blue) and dynLED condition (orange) for each participant. DLMO during the baseline night was significantly earlier than during the treatment night in sLED condition (t = 4.85, *p* = 0.008). In the dynLED condition such a delay between baseline and treatment night was not present.

### 2.4. Sleep

#### 2.4.1. Sleep Stages and Sleep Latency

Overall, no significant main effect for the factor ‘light’ was found for any of the sleep stages (i.e. N1, N2, N3, N4, REM). The factor ‘night’ indicated a tendency (F_1,13_ = 4.21, *p* = 0.06) for an increase in REMS from the baseline to the treatment night. N2 and NREMS decreased significantly from baseline to treatment night (F_1,13_ = 5.45, *p* = 0.036) and (F_1,13_ = 5.35, *p* = 0.038) respectively. N3 and N4 were not significantly different between the nights. There was no significant interaction of the factors ‘light’ and ‘night’.

##### Sleep latency to N2

There was a significant main effect of the factor ‘light’ on sleep latency to N2 (F_1,13_ = 5.76, *p* = 0.032). The factor ‘night’ yielded no significance (F_1,13_ = 1.57, *p* = 0.232). There was no significant interaction of the factors ‘light’ and ‘night’ (F_1,13_ = 0.01, *p* = 0.929) for sleep latency to N2. Since the distribution of sleep latency to N2 was not normal, we performed a non-parametric test (the Wilcoxon signed-rank test) for each night separately. During the treatment night it took participants significantly less time to fall asleep to N2 under dynLED (13.7 minutes) than under sLED (17.4 minutes, Z=2.13, *p* = 0.027; Figure 4). During the baseline night before dynLED one participant fell asleep to N2 before lights off. During the treatment night of dynLED two participants fell asleep before lights off, whereas during sLED only one participant fell asleep to N2 before lights off.

**Figure 4:**
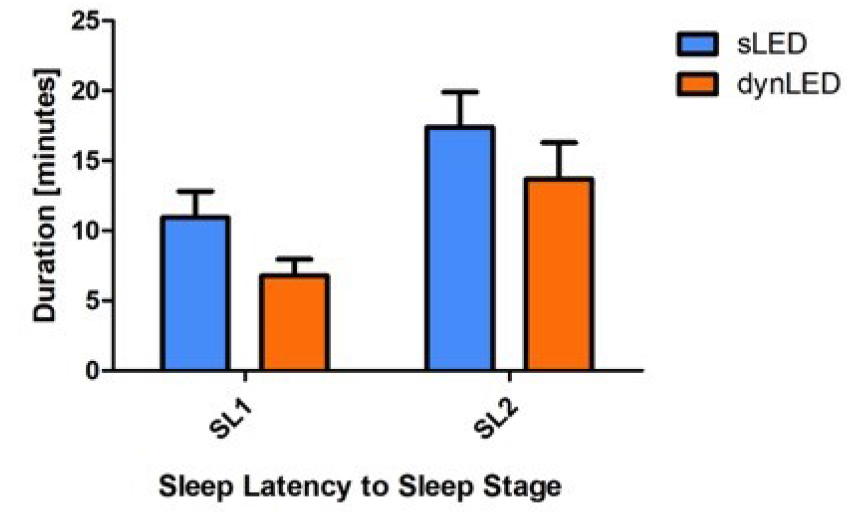
Sleep latencies to sleep stages N1 and N2 in minutes after lights off during the treatment night. Depicted are the means and standard errors of the mean for sLED (blue) and dynLED condition (orange) (n=14).

##### Sleep latency to N1

There was a significant main effect of the factor ‘light’ for sleep latency to N1 (F_1,13_ = 5.01, *p* = 0.043). The factor ‘night’ yielded no significance for sleep latency to N1 (F_1,13_ = 1.75, *p* = 0.21). There was no significant interaction of the factors ‘light’ and ‘night’ (F_1,13_ = 0.51, *p* = 0.487) for sleep latency to N1. Since the distribution of sleep latency to N1 was not normal, we performed the Wilcoxon signed-rank test for each night separately. During the treatment night participants fell significantly faster asleep to N1 under dynLED (6.8 minutes) than under sLED (10.9 minutes, Z=2.74, *p* = 0.003; Figure 4). Two participants fell asleep before lights off during the baseline night before sLED. During the baseline night before dynLED one participant the fell asleep to N1 before lights off. During the treatment night of dynLED two participants fell asleep before lights off, whereas during sLED only one participant fell asleep before lights off.

#### 2.4.2. EEG Delta Activity

EEG delta activity (0.75 – 4.5 Hz) during NREMS, when expressed relative to the baseline power density per participant in the frontal EEG derivations showed no significant effect of the factor ‘light’. The temporal dynamics of relative frontal EEG delta activity exhibited the usual decline across the night with a superimposed ultradian NREM-REMS cycling during the baseline nights and both treatment nights, sLED and dynLED (Figure 5, F_51,541_ = 16.57, *p* < 0.0001). There was a significant interaction ‘light’ x ‘time of day’ (F_50,540_ = 1.56, *p* < 0.01), but *post-hoc* comparisons indicated no significant difference between the two light conditions.

**Figure 5:**
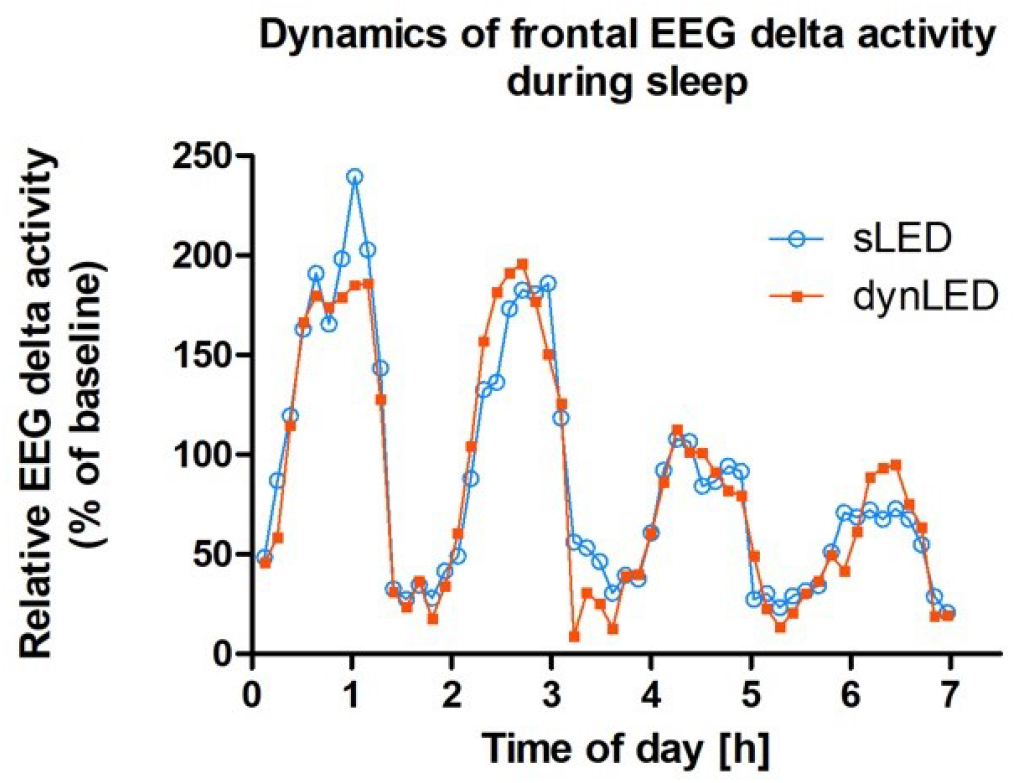
Temporal dynamics of relative EEG delta activity (0.75 - 4.5 Hz) in NREMS during the treatment night for sLED (blue) and dynLED (orange) expressed relative to the baseline power density per participant in the frontal EEG derivations. The main effect of the factor ‘light’ was not significant. Time of day is the average clock time according to the participant’s habitual bedtimes (mean values, n=14).

## 3. Discussion

The implementation of a dynamic lighting condition during scheduled wakefulness across a 16-h waking day resulted in significantly less melatonin suppression, lower subjective vigilance in the evening hours prior sleep and faster sleep onset in the following sleep episode in comparison to the same but static lighting condition. Sleep structure, sleep quality, subjective sleepiness, cognitive performance and visual comfort did not significantly differ between the two lighting conditions.

While we aimed at deploying a “naturalistic” sinusoidal illuminance change, the continuous change of CCT did not simulate a “natural daylight change”, but took into consideration the general recommendation of using blue-depleted light in the evening. However, our study design does not allow to answer the question, if the change in illuminance itself or the change in CCT itself would have led to the same observed effects. It addresses the question if a continuous change is more efficient than a square wave on/off light signal during dawn and dusk times. We deployed this continuous change according to the recommendation not to exceed the maximum speed of CCT variation of 12 K/s ^33^.

### Cognitive Performance and Sleepiness/Vigilance

In the present study, we did not find significant differences in cognitive performance between the lighting conditions. 1.000 K difference (melanopsin weighted irradiance being 24% higher under dynLED, +1.7 μW/cm^2^) during the day did not change cognitive performance although several studies reported light-induced alerting effects during daytime ^34^ (5000 lux, melanopsin weighted irradiance 373 μW/cm^2^) particularly with bright light (1000 lux, assumed melanopic weighted irradiance of 85 μW/cm^2^) ^35^ or of short-wavelength ^36^ (460 nm, 5 lux, melanopsin weighted irradiance 8,5 μW/cm^2^). In one study with 94 office workers working four weeks under blue-enriched white light (17.000 K) during the day, subjective attention, positive mood, productivity and concentration were significantly higher than under neutral white light (4.000 K) ^37^. Illuminance at eye level is not available for this study. Furthermore even an exposure of only 18 minutes during daytime to blue light with a wavelength of 470 nm can stimulate brain regions responsible for perception, memory and emotions compared to green light (550 nm)^38^. Observed improvements are probably due to the sensitivity of ipRGCs, as magnetic resonance imaging showed that blue light activates brain regions (prefrontal cortex and thalamus, responsible for alertness and cognition) even in blind individuals with still intact ganglion cells ^39^. Therefore, we hypothesized better daytime cognitive performance although other studies also reported contradictory results ^40–55^. It should be noted, that due to the high spectral quality (resulting in a very high CRI) in both lighting conditions, the sLED melanopsin weighted irradiance of 7 μW/cm^2^ was already 23% more compared to 5.7 μW/cm^2^ if a conventional white LED with a lower CRI at the same illuminance and CCT would have been used. Furthermore, the volunteers overall cognitive performance was rather high, making it difficult to further increase it by an environmental factor such as light (i.e. ceiling effect) during daytime when the circadian pacemaker fully promotes wakefulness in humans ^56^. Subjective ratings of wakefulness on a 5-point Likert scale only differed when lighting conditions were notably different, i.e. at 11 pm. As hypothesized, during the late evening participants felt rather tired under a very low illuminance (approx. 12 lux) and low CCT (approx. 2.500 K). Nevertheless, this difference was neither expressed in KSS sleepiness ratings nor cognitive performance. Notably, questionnaires were translated to German. In German there is a difference between tiredness and sleepiness meaning that although participants were (physically or cognitively) tired they were not sleepy. During the late evening, participants experienced dynamic light as being too dark. However, during the day, the 1.000 K increase in the dynLED compared to the sLED was not large enough to be visually noticeable, maybe due to adaptation (white balance) of our perception. A tendency (*p* = 0.07) for better CCT ratings during the dynamic LED condition was most prominent at 11 pm.

### Circadian Melatonin Profiles

We found 36% more attenuation of melatonin during sLED in the last sample before sleep compared to dynLED, which corroborates the results by Chellappa et al. 2011 ^57^ and fits the model proposed by Prayag et al. 2019 ^58^ well. Their findings support the assumption that melatonin suppression by light is predominantly driven by melanopsin and that it can be initiated already at low irradiances. They calculated an initiation threshold for the melatonin suppression response to light at 0.2 μW/cm^2^ melanopsin weighted irradiance. In the present study melanopsin weighted irradiance during dynLED was below this level at 0.06 μW/cm^2^ (horizontal in bed, 10 minutes prior bedtime). Melanopsin weighted irradiances one hour prior bedtime were 15 fold lower during dynLED than during sLED and 150 fold lower 10 minutes prior bedtime respectively. Consequently, we expected that melatonin suppression should be minimal during this time period in the dynLED condition.

In the review by Prayag ^58^ the saturation of melatonin suppression was assumed at 36.6 μW/cm^2^ melanopsin weighted irradiance, and the relative melatonin suppression of 50% was calculated to be at 2.5 μW/cm^2^ melanopsin weighted irradiance. During our sLED condition, melanopsin weighted irradiance was at 9.14 μW/cm^2^ with 36% stronger melatonin attenuation compared to dynLED. The reason for less melatonin suppression at higher melanopsin weighted irradiances in the present study compared to the model by Prayag ^58^ may be due to pupil constriction (miosis) during sLED. Since Prayag ^58^ analyzed a dataset from ^13^ in which pupils where dilated and volunteers were dark-adapted, lower light levels might have caused retinal irradiances to be higher than in our study. Unfortunately, retinal irradiance cannot be estimated from irradiance measured at the cornea because of the imaging optics of the eye ^59^. Furthermore, the calculation of melatonin suppression by Prayag ^58^ was control-adjusted against complete darkness whereas here we compare two lighting conditions, one of which with continuously decreasing melanopsin weighted irradiances.

It is still unclear if melanopsin in the ipRGCs works independently at all light intensities. It seems that at certain ambient brightness levels they do. Only when brightness is too low, they receive additional input from rods ^60^. Previous findings in humans suggest, melanopsin in the ipRGCs to be the primary circadian photopigment in response to long-duration light exposure and at irradiances in the photopic range ^61^. Also, more recent research suggests that the suppression of the hormone melatonin is mainly driven by the response of ipRGCs to light ^62,63^. The present study underpins, that exposure to white polychromatic light at 4000 K compared to warm white light at 2700 K and lower irradiances attenuates the secretion of melatonin in the late evening but it does not explain the mechanisms behind, because both, melanopsin and s-cone weighted irradiances, were changed simultaneously. The attenuated evening rise in melatonin did not induce a phase shift since the timing of the melatonin offset in the morning after the treatment night was not shifted. This may be due to the fact that a possible phase delay by 4000 K light (sLED) in the late evening was counteracted by a phase advance in the early morning when volunteers were woken up with 4000 K light (sLED) in comparison to a much darker light during the dynamic condition. This gives evidence for a continuous entrainment process of the circadian oscillator constantly accelerating and decelerating according to the lighting condition in the environment.

### Sleep

Due to the short sleep latencies in good sleepers we selected for our study, we did not expect differences in sleep latencies due to possible floor effects. Although there was a significant effect of the main factor ‘light’, indicating that it took participants longer to fall asleep to N2 during sLED the interaction term ‘light’ x ‘night’ was not significant. However, non-parametric testing for each night separately, yielded significant shorter sleep latencies to both N1 and N2 in the treatment night for the dynLED than the sLED condition. Since participants were already lying in bed and waiting for the lights to be switched off, it happened, that participants fell asleep before lights off.

We hypothesized more EEG delta activity based on previous findings reporting more slow-wave activity after lower CCTs in the evening ^64^. Chellappa et al. ^64^ investigated the effects of 2h/ 40 lux compact fluorescent light of various CCTs on sleep. After cold light (6.500 K) EEG slow-wave activity was reduced significantly during the first sleep cycle compared to light at 2500 K and 3000 K. Since we compared 2.700 K with 4.000 K, a difference of 1.300 K might not have been enough to elicit sleep alterations. Since we did not measure pupil size, it is not clear whether the continuously changing illuminance caused a continuous adaptation of the participants (the darker the environment, the larger the pupil size). This could have led to similar retinal illuminances under both conditions even when the environment was significantly darker during the dynLED condition. Furthermore, under low CCTs and illuminances already starting in the late afternoon in our experiment, the build up of homeostatic sleep pressure may have been attenuated compared to sLED condition with a constant CCT of 4000 K and constant illuminance. Indeed, a recent study suggests that not only the duration of prior wakefulness, but also the experienced illuminance during wakefulness affects homeostatic sleep regulation in humans ^65^. One could therefore recommend that warm light and low illuminances should be administered in the late evening, only for a very few hours before usual bedtime. Sleep latency did not change in the previous studies mentioned above (in which no dynamics were deployed). Therefore, one could assume that shorter sleep latencies as found in the present study was solely caused by the dynamics rather than the dimmer and warmer light.

## 4. Conclusion

Previous research has predominantly focused on non-visual effects of light during the night, less often during the day, and occasionally during twilight but not the combination of the three. To our best knowledge, the continuous change of lighting during 16 hours of wakefulness on human circadian physiology, cognitive performance and sleep has not been investigated so far under controlled laboratory conditions.

Our results did not confirm our hypothesis that CCT and intensity of light influences daytime cognitive performance, even with low CCT and intensity during the evening. Furthermore, participants did not report significant differences in their subjectively perceived sleepiness levels, except for the late evening hours, when CCTs and illuminances were very distinct from each other. At this time point participants reported a decrease in vigilance and perceived the room as too dark during the dynamic condition, despite this, cognitive performance did not deteriorate. The light conditions did not influence sleep architecture and subjective sleep quality.

Our study has some limitations. Since our experiment does not answer the question, if the change in illuminance or the change in CCT *per se* would have led to the same effects, this needs to be addressed in future studies. Furthermore, we still do not know if a continuous change is more efficient than a stepwise reduction of CCT and illuminance in humans.

Our findings are consistent with previous work showing that melatonin suppression was significantly enhanced at colder CCTs and higher illuminances in the evening (static condition). This can be avoided by using dynamically changing light as implemented in our study. As melatonin is important for many physiological processes in the human body (e.g. antioxidant and regulating sleep-wake timing), it should not be suppressed in the evening and night by light. Therefore, our results foster the common recommendation of using blue-depleted light and low illuminances in the late evening. During daytime we could not find any significant differences between dynamic and static light. So-called Human Centric Lighting (HCL) incorporating dynamically changing correlated color temperature (CCT) and illuminance across the day can be recommend to be used when applied during the late evening at home or during late shift work. During common day shift work the use HCL as tested in our experiment seems not to have any superior effect. While we deployed a naturalistic sinusoidal illuminance change, the continuous change of CCT did not simulate a natural change during twilight. This could be a topic for future studies.

## 5. Methods

The design and method of the present study is based on our previously published study ^66^ and thus summarized here. In contrast to the previous study comparing conventional LED spectra with enhanced LED spectra (“Daylight LED”), here we used Daylight LEDs for both, the dynamic and static light condition. The study procedures were approved by the local ethics committee and performed according to the Declaration of Helsinki. All study participants provided written informed consent.

### 5.1. Study Design

The study was carried out at the Centre for Chronobiology in Basel, Switzerland, between April 2017 and March 2018. The ‘in-laboratory part of the study’ comprised two 49-hour episodes, which participants spent in sound-attenuated chronobiology suites under light, temperature and humidity controlled conditions without any time cues (Figure 6). Volunteers reported to the laboratory 6 hours prior to their usual bedtime, when electrodes for polysomnographic recordings (PSG) were attached, cognitive test batteries explained and practiced during the first evening under standard fluorescent lighting conditions (Philips Master TL5 HO 54W/830, CRI 80, 3.000 K, resulting in 2800 K and 100 lux at the participant’s eye level in bed facing the opposite wall). After an 8-hour sleep episode baseline night, scheduled at their habitual bedtimes, volunteers either woke up in the sLED or the dynLED condition (Toshiba TRI-R Circadian System NP10576, based on TRI-R LED SMD5056) and spent 16 hours awake in one of the conditions, followed by a second 8-hour sleep episode (i.e. treatment night) and a final 11-hour episode of scheduled wakefulness. During scheduled wakefulness volunteers were allowed to move freely in their room when they were not involved in scheduled tasks. They were allowed to read and listen to music but were not allowed to use electronic devices such as mobile phones and tablet PCs. They received the same scheduled meals (25 minutes, 4 hours and 11 hours after wake up). The order of the lighting conditions (sLED and dynLED) was counterbalanced, such that half of the participants started with sLED and vice versa. The washout period between the two in-lab sessions was one week. The study protocol is shown in Figure 7.

**Figure 6:**
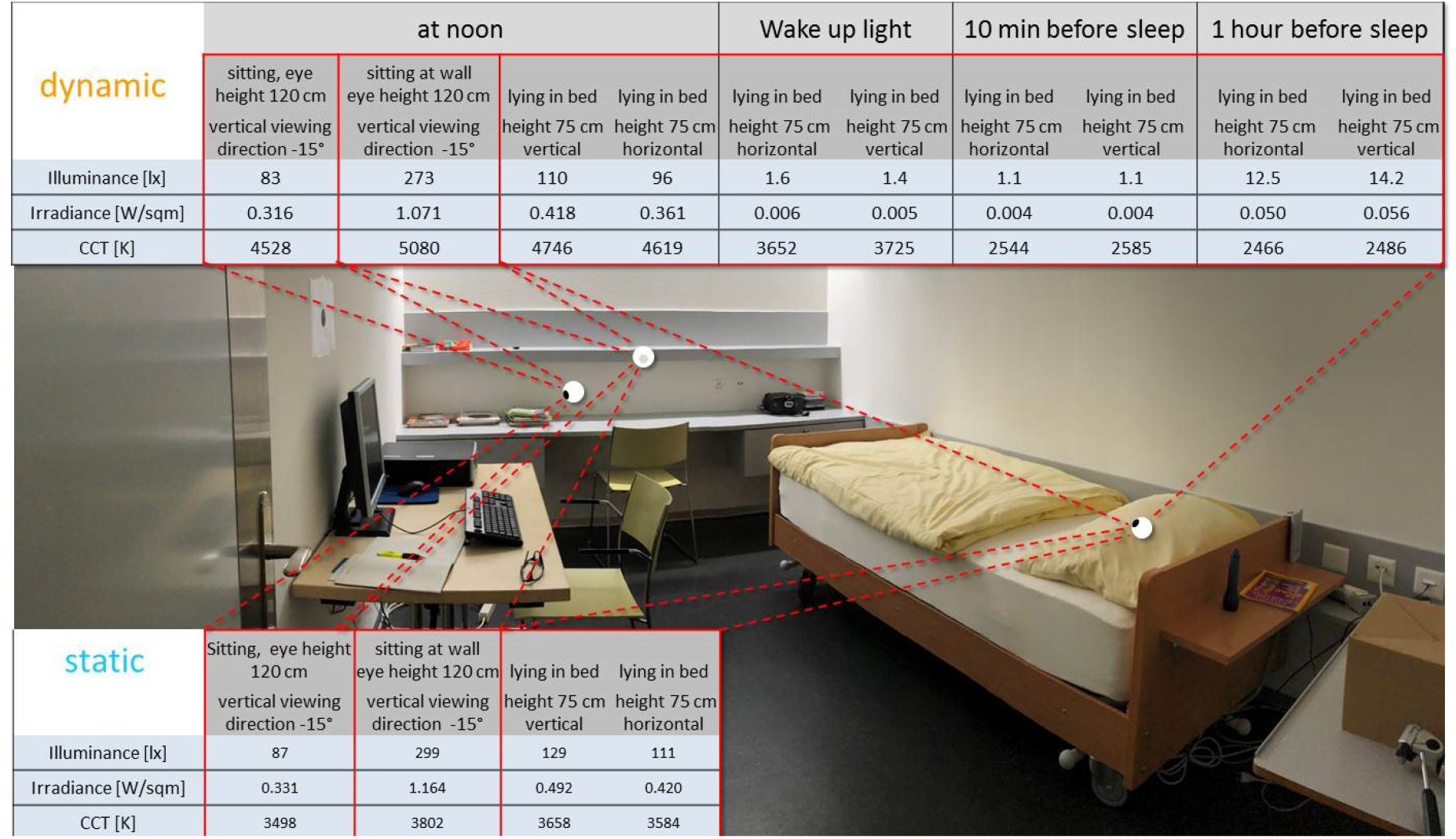
Lighting parameters during the study at different times (only for the dynamic condition) and at different locations in the laboratory for both conditions.

**Figure 7:**
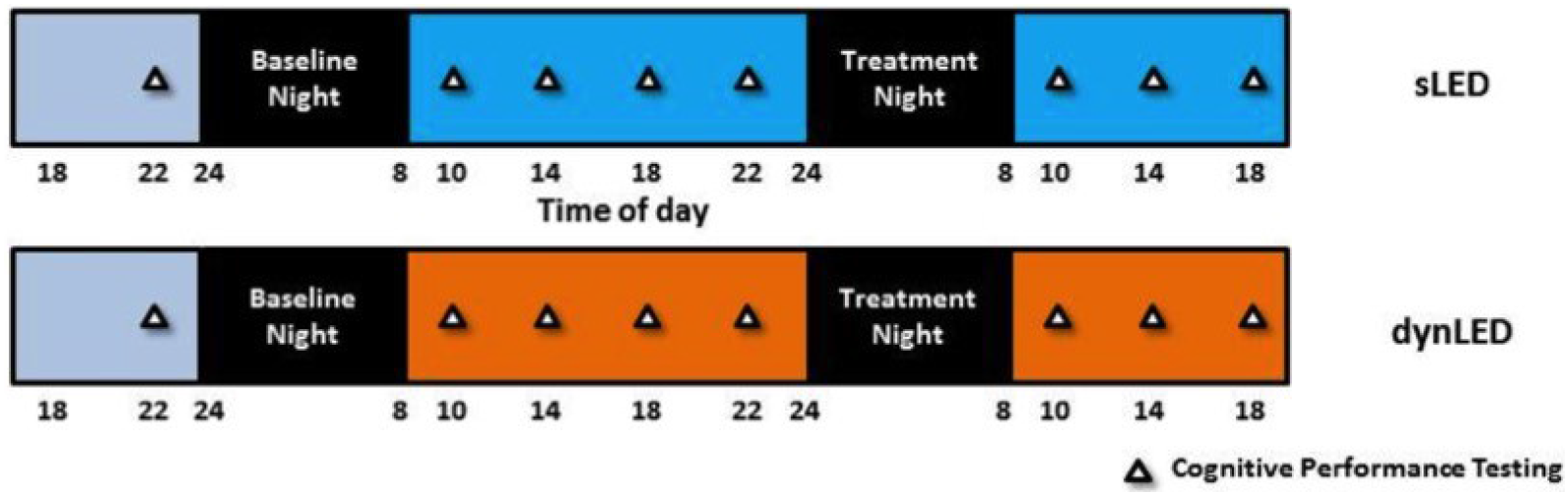
Schedule of the study: Participants spent twice 49 hours in the laboratory, once under a static LED light condition (sLED in blue) and once under dynamically changing LED light (dynLED in orange). The first evening (grey) they spent in identical conditions. Triangles show the timing of cognitive performance testing. Salivary melatonin samples were taken half hourly in the evening and every two hours during the day.

### 5.2. Participants

Eighteen healthy male participants were screened for sleep and psychiatric disorders and spent one PSG-assessed habituation night in the sleep lab prior to study participation to exclude potential sleep disorders (i.e. sleep apnea, periodic leg movements etc.). All volunteers received monetary compensation for their participation in the study. All study participants had a good sleep quality as assessed with the Pittsburgh Sleep Quality Index ^67^ (Global PSQI score ≤ 5) and were no extreme chronotypes (42 and 57 points on the Munich Chronotype questionnaire ^68^). They underwent a medical examination carried out by the physician in charge and an ophthalmic examination by a certified optometrist to exclude volunteers with visual impairments. Participants were not excluded if they wore glasses or contact lenses. All participants were screened for color deficiency by the Ishihara Test ^69^. Exclusion criteria were smoking, medication or drug consumption, shift work within the last three months and transmeridian flights up to one month prior to the study. One week prior to the laboratory admission, participants were instructed to keep a regular sleep-wake schedule (no naps, ± 60 min of habitual bedtimes) and compliance was verified by sleep logs and continuous wrist actimetry (Actiwatch, Cambridge Neurotechnology, Cambridge, UK). Additionally, participants were asked to refrain from alcohol, and a toxicological screen was performed upon laboratory entry. After dropouts due to headache (N=1), and technical issues (N=3), data of 14 participants (mean age 25.58 ± 3.34 years, intermediate chronotypes) remained for further analysis.

### 5.3. Light Treatment

During scheduled wakefulness, light exposure in the study room during the static condition was set to 4.000 K and 100 lux (vertical at the eye when sitting at the desk or lying in bed) (Figure 6). The dynamic condition was defined as a continuous light change starting in the morning with 3500 K / < 1 lux incrementally increasing until reaching a peak of approximately 5000 K / 100 lux (at the eye) during the day lasting until 3 pm. The luminance at the front wall was set to 66.7 cd/m^2^, at the left wall to 24.2 cd/m^2^ for both conditions between 10 am and 3 pm. The lighting conditions are shown in Figure 6 and Figure 8–Figure 10. Afterwards, CCT and illuminance slowly and continuously decreased in the afternoon, finally reaching 2700 K / < 1 lux at bedtime. During dynLED, the luminance during the late evening (at 23:20 h) on the pillow was 0.3cd/m^2^, meaning mesopic vision was active (both rods and cones contribute to vision). Horizontal irradiances on the pillow at that time during sLED were 100fold higher than during dynLED (42 μW/cm^2^ vs. 0.43 μW/cm^2^). Since there is no single action spectrum for NIF responses to light and a description of optical radiation solely according to the photopic action spectrum is not sufficient, we report the responses of all photoreceptors according to the new CIE S 026/E:2018 standard ^70^ (Figure 10). At 23:20 h melanopsin weighted irradiance during dynLED was 0.06 μW/cm^2^ (horizontal in bed) vs. 9.14 μW/cm^2^ for sLED. At 22:30 h (one hour prior bedtime) melanopsin weighted irradiance was 0.6 μW/cm^2^ during dynLED. **Figure 8** shows the change of melanopic irradiances across the day. The spectral characteristics are shown in Figure 9.

**Figure 8:**
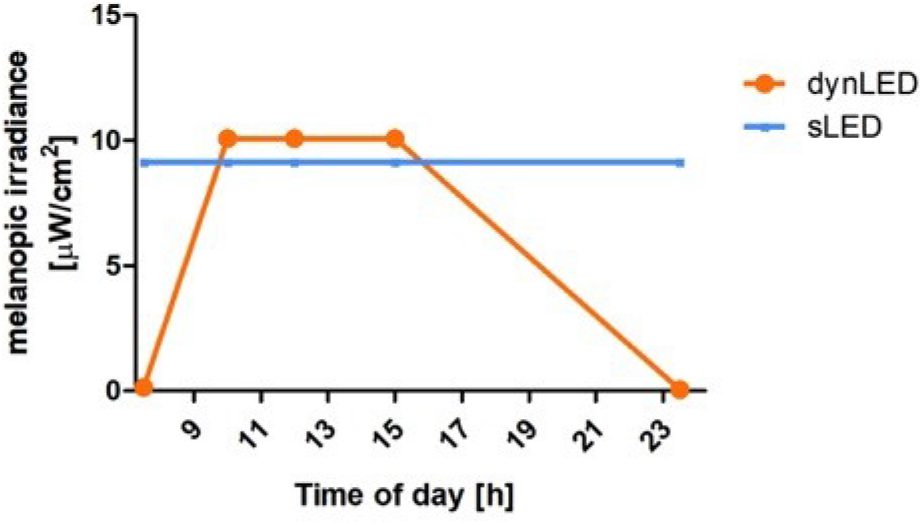
Change of melanopic irradiance [in μW/cm^2^] during the study at different times of day. Between 10 am and 4 pm melanopic irradiance is higher in the dynLED condition compared to sLED. After 4 pm and especially in the late evening melanopic irradiance is significantly lower in the dynLED condition compared to sLED.

**Figure 9:**
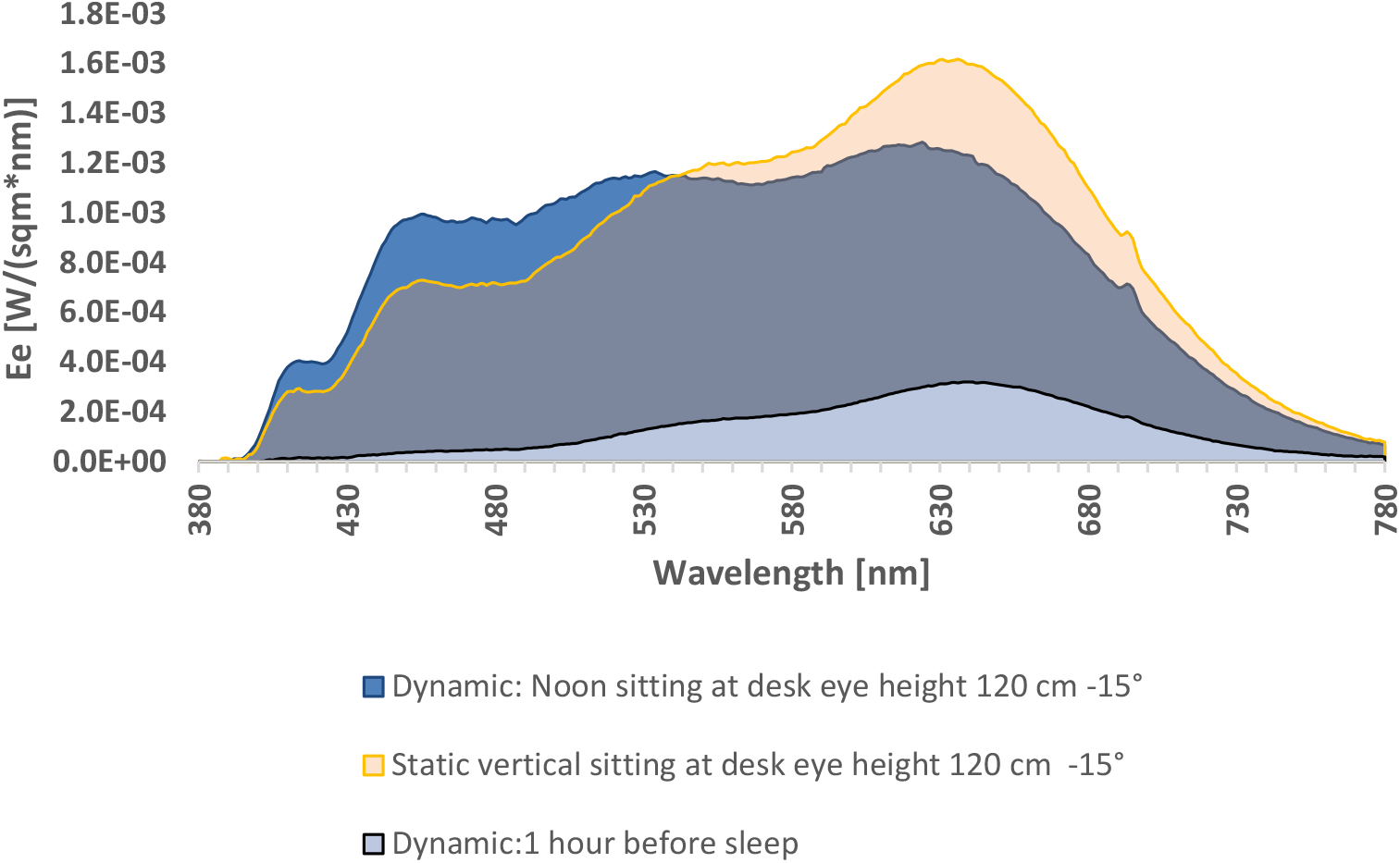
Light spectra used during the study: Spectral irradiance vertical at eye height (120 cm −15°) sitting at the desk during dynLED (between 10 am and 3 pm) and sLED (at all times). The lowest curve shows the spectral irradiance during dynLED one hour prior bed time.

**Figure 10:**
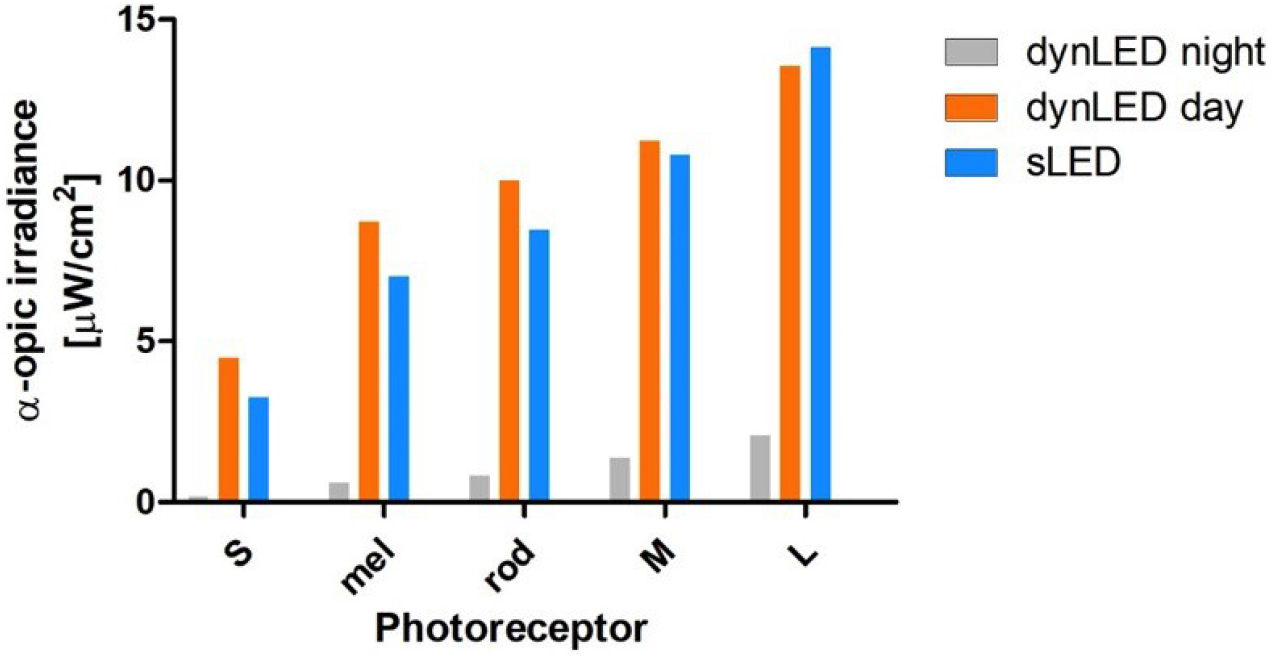
Responses of all photoreceptors according to the new CIE standard ^70^. Depicted are photoreceptor weighted irradiances: S-cones, melanopsin, rods, M-cones and L-cones during the day for sLED and dynLED and one hour prior sleep only during dynLED.

### 5.4. Cognitive Performance and Subjective Variables

During the 16 hours of scheduled wakefulness starting 70 minutes after lights on in the morning cognitive performance was assessed four times (every four hours). Participants did all tests in front of a grey computer screen. Among various subjective variables, the 35-minute cognitive test battery included a visual verbal n-back task and an auditory sustained attention task (i.e. psychomotor vigilance task, PVT).

#### 5.4.1. Working Memory Performance (n-back paradigm)

During the n-back task volunteers indicated if a displayed letter matches a target stimulus that was presented n trials ago. White letters were used on a grey screen. Each session consisted of six blocks, divided into two bouts. The demand level was adjusted to individual performance of the volunteers. The participants were able to familiarize themselves with the n-back level during a brief practice period. For further information about the n-back task please refer to ^66^.

#### 5.4.2. Psychomotor Vigilance Performance Task

The psychomotor vigilance performance task is a sustained attention task, that is sensitive to circadian rhythmicity and sleep need ^71^. Volunteers were requested to press a response button as fast as possible as soon they heard an auditory stimulus. They should also avoid pressing the button too soon. The task lasted 10 minutes during which the stimulus was presented in intervals randomly varying from 2 to 9 seconds.

#### 5.4.3. Subjective Sleepiness and Visual Comfort

During the entire laboratory stay, volunteers periodically rated their sleepiness levels on the Karolinska Sleepiness Scale (KSS) ^72^ as follows: Immediately after wake up until three hours after and in the evening starting five hours prior to lights off in hourly or two hourly intervals. To assess volunteer’s subjective perception of visual comfort, we used a five-point Likert type scale that probed brightness, and correlated color temperature based on a selection of questions derived from Eklund and Boyce ^73^.

### 5.5. Melatonin

Saliva collections were scheduled every 30 minutes in the morning and evening and every 1 or 2 hours in between (for precise timing see Figure 2). A direct double-antibody radioimmunoassay was used for the melatonin assay (validated by GC-MS with an analytical least detectable dose of 0.65 pg/ml; Bühlmann Laboratory, Schönenbuch, Switzerland). The minimum detectable dose of melatonin (analytical sensitivity) was determined to be 0.2 pg/ml.

In order to assess changes in circadian phase, dim light melatonin onset (DLMO) was calculated for the baseline and treatment evening. We fitted the evening melatonin profile by a piecewise linear-parabolic function using the interactive computer-based hockey-stick algorithm to calculate the DLMO for each participant ^74^. An ascending level of 2.5 was applied; however, we used an ascending level of 1.5 for five evenings when melatonin concentration was too low.

### 5.6. PSG

Sleep EEG activity was continuously recorded during sleep with the Vitaport Ambulatory system (Vitaport-3 digital recorder TEMEC Instruments BV, Kerkrade, the Netherlands). Twelve EEG derivations (Fz, F3, F4, Cz, C3, C4, Pz, P3, P4, Oz, O1, O2) referenced against linked mastoids (A1 and A2), two electrooculograms, one submental electromyogram and one electrocardiogram were recorded. All signals were low pass filtered at 30Hz at a time constant of 1 second.

Sleep stages were visually scored per 30-second epochs according the standard criteria ^75^. Non-rapid eye movement sleep (NREMS) was defined as the sum of NREM stages 2, 3 and 4. Slow wave sleep (SWS) was defined as the sum of NREMS stages 3 and 4. Spectral analysis was conducted using a fast Fourier transformation, which produced a 0.25Hz bin resolution. EEG power spectra were calculated during NREMS in the frequency range from 0 to 32Hz. Artefact-free 4-second epochs were averaged across 30-second epochs. Here we report EEG data for frontal (Fz, F3, F4) derivations, in the frequency range of 0.75-20 Hz.

### 5.7. Statistical Analysis

Statistical analyses were performed using SAS (version 9.4; SAS Institute, Cary, NC). An alpha level of 0.05 was used to assess statistical significance. All output variables of the 14 men were statistically analyzed with mixed-model analyses of variance (PROC MIXED) with main repeated factors being ‘light’ (dynLED and sLED) and ‘time of day’ and volunteers as a random factor. For PVT performance, the default performance metrics – median reaction time (RT), 10% fastest and 10% slowest RT and lapses were calculated according to Blatter et al. ^71^. Response times below 100 ms were considered as false starts and therefore excluded. For n-back performance, the following metrics were used: number of hits, false alarms, accuracy and the percentage of correct responses. All-night EEG power density in NREMS was analyzed for frontal derivations for each 0.25 Hz frequency bin, with the main factor ‘light’. NREM-REMS cycles were defined according to an adapted method from Feinberg and Floyd ^76^. Thereof, each sleep cycle was subdivided into ten time intervals of equal length during NREMS and into four time intervals during REMS.

## Contributions

OS and CC wrote the main manuscript text. CC and YS designed the experiments. OS, MF, SV, TB and JW carried out the study and were responsible for the study administration. OS, and CC were responsible for the analysis of the data. CC served as supervisor of the study. MM was one of the study doctors. All authors provided critical review of and revisions to the manuscript. All authors have approved the final version of this manuscript.

## Acknowledgements

We thank all the volunteers for participating in the study. We also thank all the study doctors and study helpers for their assistance during the study, especially the night shifts.

## Data availability

The datasets analyzed during this study are available from the corresponding author upon reasonable request.

## REFERENCES

1 Schweizer, C. et al. Indoor time-microenvironment-activity patterns in seven regions of Europe. Journal of Exposure Science & Environmental Epidemiology 17, 170–181, doi:10.1038/sj.jes.7500490 (2007).

2 Ohayon, M. M. & Milesi, C. Artificial Outdoor Nighttime Lights Associate with Altered Sleep Behavior in the American General Population. Sleep 39, 1311–1320, doi:10.5665/sleep.5860 (2016).

3 Ohayon, M. et al. National Sleep Foundation’s sleep quality recommendations: first report. Sleep Health 3, 6–19, doi:10.1016/j.sleh.2016.11.006 (2017).

4 Kozaki, T., Miura, N., Takahashi, M. & Yasukouchi, A. Effect of reduced illumination on insomnia in office workers. J Occup Health 54, 331–335, doi:10.1539/joh.12-0049-fs (2012).

5 Mason, I. C. et al. Circadian Health and Light: A Report on the National Heart, Lung, and Blood Institute’s Workshop. J Biol Rhythms 33, 451–457, doi:10.1177/0748730418789506 (2018).

6 Daneault, V., Dumont, M., Masse, E., Vandewalle, G. & Carrier, J. Light-sensitive brain pathways and aging. J Physiol Anthropol 35, 9, doi:10.1186/s40101-016-0091-9 (2016).

7 Benarroch, E. E. The melanopsin system: Phototransduction, projections, functions, and clinical implications. Neurology 76, 1422–1427, doi:10.1212/WNL.0b013e31821671a5 (2011).

8 LeGates, T. A., Fernandez, D. C. & Hattar, S. Light as a central modulator of circadian rhythms, sleep and affect. Nat Rev Neurosci 15, 443–454, doi:10.1038/nrn3743 (2014).

9 Gaggioni, G., Maquet, P., Schmidt, C., Dijk, D. J. & Vandewalle, G. Neuroimaging, cognition, light and circadian rhythms. Front Syst Neurosci 8, 126, doi:10.3389/fnsys.2014.00126 (2014).

10 Hattar, S., Liao, H. W., Takao, M., Berson, D. M. & Yau, K. W. Melanopsin-containing retinal ganglion cells: architecture, projections, and intrinsic photosensitivity. Science 295, 1065–1070, doi:10.1126/science.1069609 (2002).

11 Dacey, D. M. et al. Melanopsin-expressing ganglion cells in primate retina signal colour and irradiance and project to the LGN. Nature 433, 749–754, doi:10.1038/nature03387 (2005).

12 Thapan, K., Arendt, J. & Skene, D. J. An action spectrum for melatonin suppression: evidence for a novel non-rod, non-cone photoreceptor system in humans. J Physiol 535, 261–267, doi:10.1111/j.1469-7793.2001.t01-1-00261.x (2001).

13 Brainard, G. C. et al. Action spectrum for melatonin regulation in humans: evidence for a novel circadian photoreceptor. J Neurosci 216405–6412 (2001).

14 Benloucif, S. et al. Stability of melatonin and temperature as circadian phase markers and their relation to sleep times in humans. J Biol Rhythms 20, 178–188, doi:10.1177/0748730404273983 (2005).

15 Cajochen, C. et al. High sensitivity of human melatonin, alertness, thermoregulation, and heart rate to short wavelength light. J Clin Endocrinol Metab 90, 1311–1316, doi:10.1210/jc.2004-0957 (2005).

16 Minors, D. S., Waterhouse, J. M. & Wirz-Justice, A. A human phase-response curve to light. Neuroscience Letters 133, 36–40, doi:https://doi.org/10.1016/0304-3940(91)90051-T (1991).

17 Khalsa, S. B., Jewett, M., Cajochen, C. & Czeisler, C. A Phase Response Curve to Single Bright Light Pulses in Human Subjects. The Journal of physiology 549, 945–952, doi:10.1113/jphysiol.2003.040477 (2003).

18 Ruger, M. et al. Human phase response curve to a single 6.5 h pulse of short-wavelength light. J Physiol 591, 353–363, doi:10.1113/jphysiol.2012.239046 (2013).

19 Gooley, J. J. et al. Exposure to room light before bedtime suppresses melatonin onset and shortens melatonin duration in humans. J Clin Endocrinol Metab 96, E463–472, doi:10.1210/jc.2010-2098 (2011).

20 Burgess, H. J. & Molina, T. A. Home lighting before usual bedtime impacts circadian timing: a field study. Photochem Photobiol 90, 723–726 (2014).

21 Boulos, Z., Macchi, M. & Terman, M. Twilight transitions promote circadian entrainment to lengthening light-dark cycles. Am J Physiol 271, R813–818, doi:10.1152/ajpregu.1996.271.3.R813 (1996).

22 Boulos, Z., Macchi, M. M. & Terman, M. Twilights widen the range of photic entrainment in hamsters. J Biol Rhythms 17, 353–363, doi:10.1177/074873002129002654 (2002).

23 Boulos, Z., Macchi, M. & Terman, M. Effects of twilights on circadian entrainment patterns and reentrainment rates in squirrel monkeys. J Comp Physiol A 179, 687–694 (1996).

24 Boulos, Z., Macchi, M., Houpt, T. A. & Terman, M. Photic entrainment in hamsters: effects of simulated twilights and nest box availability. J Biol Rhythms 11, 216–233, doi:10.1177/074873049601100304 (1996).

25 Daan, S. Colin Pittendrigh, Jürgen Aschoff, and the Natural Entrainment of Circadian Systems. Journal of Biological Rhythms 15, 195–207, doi:10.1177/074873040001500301 (2000).

26 Danilenko, K. V., Wirz-Justice, A., Krauchi, K., Weber, J. M. & Terman, M. The human circadian pacemaker can see by the dawn’s early light. J Biol Rhythms 15, 437–446, doi:10.1177/074873000129001521 (2000).

27 Gabel, V. et al. Effects of artificial dawn and morning blue light on daytime cognitive performance, well-being, cortisol and melatonin levels. Chronobiol Int 30, 988–997, doi:10.3109/07420528.2013.793196 (2013).

28 Gabel, V. et al. Dawn simulation light impacts on different cognitive domains under sleep restriction. Behav Brain Res 281, 258–266, doi:10.1016/j.bbr.2014.12.043 (2015).

29 Viola, A. U. et al. Dawn simulation light: a potential cardiac events protector. Sleep Med 16, 457–461, doi:10.1016/j.sleep.2014.12.016 (2015).

30 Terman, M. & Terman, J. S. Controlled trial of naturalistic dawn simulation and negative air ionization for seasonal affective disorder. The American journal of psychiatry 163, 2126–2133, doi:10.1176/ajp.2006.163.12.2126 (2006).

31 Terman, M., Schlager, D., Fairhurst, S. & Perlman, B. Dawn and dusk simulation as a therapeutic intervention. Biol Psychiatry 25, 966–970 (1989).

32 de Kort, Y. & Smolders, K. Effects of dynamic lighting on office workers: First results of a field study with monthly alternating settings. Lighting Research & Technology 42, 345–360, doi:10.1177/1477153510378150 (2010).

33 Bieske, K. in Proceedings of CIE2010 “Lighting Quality and Energy Efficiency. 290–297.

34 Ruger, M., Gordijn, M. C., Beersma, D. G., de Vries, B. & Daan, S. Time-of-day-dependent effects of bright light exposure on human psychophysiology: comparison of daytime and nighttime exposure. Am J Physiol Regul Integr Comp Physiol 290, R1413–1420, doi:10.1152/ajpregu.00121.2005 (2006).

35 Phipps-Nelson, J., Redman, J. R., Dijk, D. J. & Rajaratnam, S. M. Daytime exposure to bright light, as compared to dim light, decreases sleepiness and improves psychomotor vigilance performance. Sleep 26, 695–700, doi:10.1093/sleep/26.6.695 (2003).

36 Rahman, S. A. et al. Diurnal spectral sensitivity of the acute alerting effects of light. Sleep 37, 271–281, doi:10.5665/sleep.3396 (2014).

37 Viola, A. U., James, L. M., Schlangen, L. J. & Dijk, D. J. Blue-enriched white light in the workplace improves self-reported alertness, performance and sleep quality. Scand J Work Environ Health 34, 297–306 (2008).

38 Vandewalle, G. et al. Wavelength-dependent modulation of brain responses to a working memory task by daytime light exposure. Cereb Cortex 17, 2788–2795, doi:10.1093/cercor/bhm007 (2007).

39 Vandewalle, G. et al. Blue light stimulates cognitive brain activity in visually blind individuals. J Cogn Neurosci 25, 2072–2085, doi:10.1162/jocn_a_00450 (2013).

40 Lok, R., Smolders, K., Beersma, D. G. M. & de Kort, Y. A. W. Light, Alertness, and Alerting Effects of White Light: A Literature Overview. J Biol Rhythms 33, 589–601, doi:10.1177/0748730418796443 (2018).

41 Badia, P., Myers, B., Boecker, M., Culpepper, J. & Harsh, J. R. Bright light effects on body temperature, alertness, EEG and behavior. Physiol Behav 50, 583–588 (1991).

42 Akerstedt, T., Landstrom, U., Bystrom, M., Nordstrom, B. & Wibom, R. Bright light as a sleepiness prophylactic: a laboratory study of subjective ratings and EEG. Percept Mot Skills 97, 811–819, doi:10.2466/pms.2003.97.3.811 (2003).

43 Borragán, G., Deliens, G., Peigneux, P. & Leproult, R. in Neuroergonomics (eds Hasan Ayaz & Frédéric Dehais) 221 (Academic Press, 2018).

44 Huiberts, L. M., Smolders, K. C. & de Kort, Y. A. Non-image forming effects of illuminance level: Exploring parallel effects on physiological arousal and task performance. Physiol Behav 164, 129–139, doi:10.1016/j.physbeh.2016.05.035 (2016).

45 Huiberts, L. M., Smolders, K. & De Kort, Y. A. W. Seasonal and time-of-day variations in acute non-image forming effects of illuminance level on performance, physiology, and subjective well-being. Chronobiol Int 34, 827–844, doi:10.1080/07420528.2017.1324471 (2017).

46 Kaida, K. et al. Indoor exposure to natural bright light prevents afternoon sleepiness. Sleep 29, 462–469, doi:10.1093/sleep/29.4.462 (2006).

47 Leichtfried, V. et al. Intense illumination in the morning hours improved mood and alertness but not mental performance. Appl Ergon 46 Pt A, 54–59, doi:10.1016/j.apergo.2014.07.001 (2015).

48 Lok, R., Woelders, T., Gordijn, M. C. M., Hut, R. A. & Beersma, D. G. M. White Light During Daytime Does Not Improve Alertness in Well-rested Individuals. J Biol Rhythms 33, 637–648, doi:10.1177/0748730418796036 (2018).

49 Maierova, L. et al. Diurnal variations of hormonal secretion, alertness and cognition in extreme chronotypes under different lighting conditions. Sci Rep 6, 33591, doi:10.1038/srep33591 (2016).

50 Sahin, L., Wood, B. M., Plitnick, B. & Figueiro, M. G. Daytime light exposure: effects on biomarkers, measures of alertness, and performance. Behav Brain Res 274, 176–185, doi:10.1016/j.bbr.2014.08.017 (2014).

51 Smolders, K. C. H. J. & de Kort, Y. A. W. Bright light and mental fatigue: Effects on alertness, vitality, performance and physiological arousal. Journal of Environmental Psychology 39, 77–91, doi:https://doi.org/10.1016/j.jenvp.2013.12.010 (2014).

52 Smolders, K. C., de Kort, Y. A. & Cluitmans, P. J. A higher illuminance induces alertness even during office hours: findings on subjective measures, task performance and heart rate measures. Physiol Behav 107, 7–16, doi:10.1016/j.physbeh.2012.04.028 (2012).

53 Smolders, K., Peeters, S. T., Vogels, I. & de Kort, Y. A. W. Investigation of Dose-Response Relationships for Effects of White Light Exposure on Correlates of Alertness and Executive Control during Regular Daytime Working Hours. J Biol Rhythms 33, 649–661, doi:10.1177/0748730418796438 (2018).

54 TeKulve, M., Schlangen, L., Schellen, L., Souman, J. L. & van Marken Lichtenbelt, W. Correlated colour temperature of morning light influences alertness and body temperature. Physiol Behav 185, 1–13, doi:10.1016/j.physbeh.2017.12.004 (2018).

55 Vandewalle, G. et al. Daytime light exposure dynamically enhances brain responses. Curr Biol 16, 1616–1621, doi:10.1016/j.cub.2006.06.031 (2006).

56 Dijk, D.-J., Duffy, J. F. & Czeisler, C. A. Circadian and sleep/wake dependent aspects of subjective alertness and cognitive performance. Journal of Sleep Research 1, 112–117, doi:10.1111/j.1365-2869.1992.tb00021.x (1992).

57 Chellappa, S. L. et al. Non-visual effects of light on melatonin, alertness and cognitive performance: can blue-enriched light keep us alert? PLoS One 6, e16429, doi:10.1371/journal.pone.0016429 (2011).

58 Prayag, A. S., Najjar, R. P. & Gronfier, C. Melatonin suppression is exquisitely sensitive to light and primarily driven by melanopsin in humans. J Pineal Res 66, e12562, doi:10.1111/jpi.12562 (2019).

59 Spitschan, M. et al. How to Report Light Exposure in Human Chronobiology and Sleep Research Experiments. Clocks Sleep 1, 280–289, doi:10.3390/clockssleep1030024 (2019).

60 Cao, Y. et al. Mechanism for Selective Synaptic Wiring of Rod Photoreceptors into the Retinal Circuitry and Its Role in Vision. Neuron 87, 1248–1260, doi:10.1016/j.neuron.2015.09.002 (2015).

61 Gooley, J. J. et al. Spectral responses of the human circadian system depend on the irradiance and duration of exposure to light. Sci Transl Med 2, 31ra33, doi:10.1126/scitranslmed.3000741 (2010).

62 Souman, J. L. et al. Spectral Tuning of White Light Allows for Strong Reduction in Melatonin Suppression without Changing Illumination Level or Color Temperature. Journal of Biological Rhythms 0, 0748730418784041, doi:10.1177/0748730418784041.

63 Allen, A. E., Hazelhoff, E. M., Martial, F. P., Cajochen, C. & Lucas, R. J. Exploiting Metamerism to Regulate the impact of a Visual Display on Alertness and Melatonin Suppression Independent of Visual Appearance. Sleep, doi:10.1093/sleep/zsy100 (2018).

64 Chellappa, S. L. et al. Acute exposure to evening blue-enriched light impacts on human sleep. Journal of sleep research 22, 573–580, doi:10.1111/jsr.12050 (2013).

65 Cajochen, C. et al. Evidence That Homeostatic Sleep Regulation Depends on Ambient Lighting Conditions during Wakefulness. Clocks & amp; Sleep 1, 517–531 (2019).

66 Cajochen, C. et al. Effect of daylight LED on visual comfort, melatonin, mood, waking performance and sleep. Lighting Research & Technology, 1477153519828419, doi:10.1177/1477153519828419 (2019).

67 Buysse, D. J., Reynolds, C. F., 3rd, Monk, T. H., Berman, S. R. & Kupfer, D. J. The Pittsburgh Sleep Quality Index: a new instrument for psychiatric practice and research. Psychiatry Res 28, 193–213, doi:10.1016/0165-1781(89)90047-4 (1989).

68 Roenneberg, T., Wirz-Justice, A. & Merrow, M. Life between Clocks: Daily Temporal Patterns of Human Chronotypes. Journal of Biological Rhythms 18, 80–90, doi:10.1177/0748730402239679 (2003).

69 Ishihara, S. Tests for color-blindness.. Handaya, Tokyo, Hongo Harukicho (1917).

70 CIE. CIE System for Metrology of Optical Radiation for ipRGC-Influenced Responses to Light, <http://www.cie.co.at/publications/cie-system-metrology-optical-radiation-iprgc-influenced-responses-light-0> (2018).

71 Blatter, K. et al. Gender and age differences in psychomotor vigilance performance under differential sleep pressure conditions. Behav Brain Res 168, 312–317, doi:10.1016/j.bbr.2005.11.018 (2006).

72 Gillberg, M., Kecklund, G. & Akerstedt, T. Relations between performance and subjective ratings of sleepiness during a night awake. Sleep 17, 236–241, doi:10.1093/sleep/17.3.236 (1994).

73 Eklund, N. H. & Boyce, P. R. The Development of a Reliable, Valid, and Simple Office Lighting Survey. Journal of the Illuminating Engineering Society 25, 25–40, doi:10.1080/00994480.1996.10748145 (1996).

74 Danilenko, K. V., Verevkin, E. G., Antyufeev, V. S., Wirz-Justice, A. & Cajochen, C. The hockey-stick method to estimate evening dim light melatonin onset (DLMO) in humans. Chronobiol Int 31, 349–355, doi:10.3109/07420528.2013.855226 (2014).

75 A manual of standardized terminology, techniques and scoring system for sleep stages of human subjects. Allan Rechtschaffen and Anthony Kales, editors. (U. S. National Institute of Neurological Diseases and Blindness, Neurological Information Network, 1968).

76 Feinberg, I. & Floyd, T. C. Systematic trends across the night in human sleep cycles. Psychophysiology 16, 283–291 (1979).

